# Distinguishing within- from between-individual effects: How to use the within-individual centering method for quadratic patterns

**DOI:** 10.1101/2020.05.25.114421

**Authors:** Rémi Fay, Julien Martin, Floriane Plard

**Affiliations:** Swiss Ornithological Institute, Seerose 1, CH–6204 Sempach; Centre for Biodiversity Dynamics, Department of Biology, Norwegian University of Science and Technology, Trondheim, Norway; Department of Biological Sciences, University of Ottawa, Ottawa, Canada; UMR CNRS 5558 Biométrie et Biologie Evolutive, Univ Lyon; Université Claude Bernard (Lyon I), Villeurbanne, France

**Keywords:** age pattern, individual trajectory, linear mixed models, selective appearance, selective disappearance, senescence

## Abstract

1. Any average pattern observed at the population level may confound two different types of processes: some processes that occur among individuals and others that occur within individuals. Separating within- from among-individual processes is critical for our understanding of ecological and evolutionary dynamics.
2. The within-individual centering method allows distinguishing within- from among-individual processes and this method has been largely used in ecology to investigate both linear and quadratic patterns. Here we show that two alternative equations could be used for the investigation of quadratic within-individual patterns. We explain under which hypotheses each is valid. Reviewing the literature, we found that mainly one of these two equations has been used by the studies investigating quadratic patterns. Yet this equation could be inappropriate in many cases.
3. We show that these two alternative equations make different assumptions about the shape of the within-individual pattern. The choice of using one equation instead of the other should depend upon the biological process investigated. One equation assumes that all individuals show the same quadratic pattern over the whole range of the explanatory variable whereas the other assumes that the quadratic patterns depend on the average explanatory variable of each individual. We give examples of biological processes corresponding to each equation.
4. Using simulations, we showed that a mismatch between the assumptions made by the equation used to analyze the data and the biological process investigated led to flawed inference affecting both output of model selection and accuracy of estimates. We stress that the equation used should be chosen carefully to ensure that the assumption made about the shape of the within-individual pattern matches the biological process investigated. We hope that this manuscript will encourage the use of the within-individual centering method, promoting its correct application for non-linear relationships.

## Introduction

The task of distinguishing trait variation occurring within individuals from that occurring among individuals is a recurrent challenge in ecology (Vaupel et al. 1979; Vaupel and Yashin 1985; Bolnick et al. 2011; Vindenes and Langangen 2015; Hamel et al. 2018). As both within- and among-individual effects drive trait variation, the patterns observed at the population level do not necessarily reflect the changes occurring at the individual level, and vice versa. A famous example is the investigation of the relationships between a response variable and individual age (van de Pol and Verhulst 2006; Nussey et al. 2008). Variation in the response variable may arise from within-individual variation, e.g., aging effect due to accumulating experience in early-life or/and due to senescence in late life, as well as from among-individual variation, by the selective disappearance of frail individuals (Aubry et al. 2009; Bouwhuis et al. 2009; Hämäläinen et al. 2014). Indeed, in heterogeneous populations, frail individuals are expected to die first, leaving robust individuals to be overrepresented at old ages (Pigeon et al. 2017; Vedder and Bouwhuis 2018). This change in the composition of the population over time generates positive age-related relationships at the population level. Thus, the average age-related trajectory observed at the population level could differ from the average trajectory observed at the individual scale, because independent processes, arising within- and among-individuals, co-occur (Vaupel and Yashin 1985).

This problem of distinguishing between within-individual and among-individual patterns is not specific to age-related patterns. It may concern the relationships between any response variable and any type of explanatory traits, i.e., behavioral, morphometric, physiological, environmental, each time several observations per individual are obtained and aggregated (e.g., Pick et al. 2016; Dammhahn et al. 2017; Morrongiello et al. 2019; Siracusa et al. 2019). Analyzing such aggregated data, we need to distinguish the within-individual effect, i.e., how variation in the explanatory variable X within an individual affects the response variable Y, from the among-individual effect, i.e., how the average value of X for a given individual is related to the average value of Y. This problem could even be extended beyond the individual level since similar problems could be encountered every time data are aggregated at different scales, a common data structure in ecology and evolution. For the rest of the manuscript, we will mainly focus on the individual level, but similar distinctions could be relevant when comparing patterns within and among clusters of data, e.g. groups of individuals, populations, species and so on (e.g., Dakin et al. 2018; Lundin et al. 2019; Lemoine et al. 2020).

From an eco-evolutionary perspective, it is crucial to separate within- from among-individual variation in traits (Bolnick et al. 2011). For example, the investigation of life-history trade-offs, which are within-individual effects, requires accounting for among-individual variation in resource acquisition, which is an among-individual effect (van Noordwijk and de Jong 1986). Similarly, our understanding of the age-related change in demographic rates has been limited for a long time because most studies considered only the population-level pattern (Forslund and Pärt 1995; Nussey et al. 2008). Patterns within and among individuals provide different information which are of interest to understand the behavior, physiology or demography of biological systems. A within-individual effect may provide information on processes involving individual plasticity such as phenotypic change, learning or the occurrence of trade-off. On the other hand, among-individual effects depict consistent inter-individual differences and thus may suggest among-individual differences in fitness, personality or pace of life, for instance (Wilson and Nussey 2010; Reid et al. 2010; Dingemanse and Dochtermann 2013). Ignoring the distinction between within- and among-individual effects may thus lead to flawed inferences confounding processes occurring at different levels (Vaupel and Yashin 1985; van de Pol and Verhulst 2006; Kendall et al. 2011; Stover et al. 2012).

A statistical method to distinguish within- from among-individual effects is the within-individual centering method (Kreft et al. 1995; Hofmann and Gavin 1998; Snijders and Bosker 1999). This powerful method is simple to use and gained popularity rapidly following its introduction to ecologists (van de Pol and Wright 2009). To date, this work has been cited 479 times and the approach has been used in 346 published studies (research among the papers citing van de Pol and Wright 2009 using Web of Science in January 2021). While this method has been presented in the context of behavioral studies for linear patterns (van de Pol and Wright 2009), it has been used in various ecological fields including demography, aging, eco-physiology, and parasitology, and has been extended to investigate quadratic patterns (Fig. 1).

**Figure 1:**
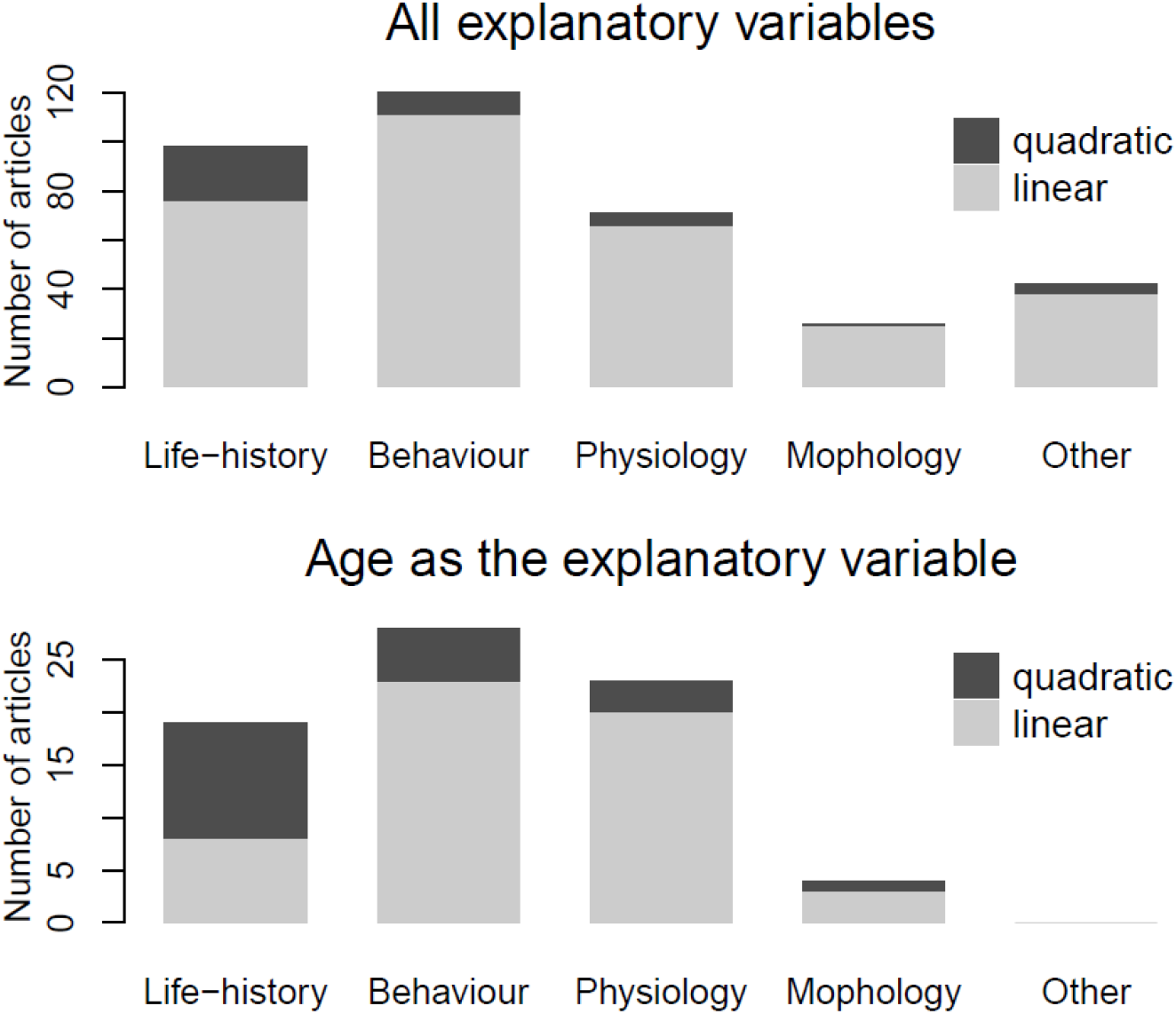
Number of articles citing the van de Pol & Wright (2009)’s work and using the within-individual centering method including all explanatory variables (upper panel) and age in particular with linear (light grey) and quadratic (dark grey) models. The articles have been classified according to the type of response variable(s) used (life-history, behavioral, physiological, morphological, and other traits).

In this paper we clarify the use of the within-individual centering method for quadratic patterns providing guidelines for its application. After briefly presenting the within-individual centering approach in case of linear patterns, we present the two alternative equations which can be used for the investigation of quadratic within-individual effects. We explain the different assumptions made by each equation about the shape of the within-individual pattern and suggest in which circumstances each equation should be used. We stress that previous studies investigating quadratic patterns failed to directly recognize these assumptions and, in some cases, may have used an inappropriate equation. Based on simulations, we assess the consequences of using the inappropriate equation on model selection and quality of estimates (bias and precision). We also illustrate the consequences of using the inappropriate equation for the specific case of quadratic-age patterns, which correspond to the most frequent individual-centered variable used when investigating quadratic individual patterns. We hope that this manuscript will encourage the use of the within-individual centering method promoting its correct application for non-linear relationships.

## 1. The within-individual centering method

In ecology, linear mixed models are nowadays standard tools to analyze data aggregated at different scales. However, such models may fail to estimate the true within-individual changes because the individual random effect may not capture the total among-individual effects (van de Pol & Wright 2009). The within-individual centering method is an extension of a simple mixed model that allows distinguishing explicitly within- from among-individual variation using additional fixed effects (Kreft et al. 1995; Hofmann and Gavin 1998; Snijders and Bosker 1999, van de Pol & Wright 2009). We present only briefly this method and refer to van de Pol & Wright (2009) for further information.

The standard random effect model can be described by the following regression equation:

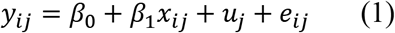

where *β_0_* is the intercept, *β_1_* is the slope, *x_ij_* is the value of a trait *x* at measurement *i* from individual *j*, *u_j_* is the deviation from the intercept for individual *j* assumed to be drawn from a normal distribution with a mean of zero and among-subject variance 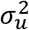, and *e_ij_* is the residual error for each measurement assumed to be drawn from a normal distribution with a mean of zero and variance 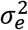.

Starting from this standard mixed model, the within-individual centering approach decomposes the effect of x in two terms: the within-individual and the among-individual effects. The among-individual effect is fitted as the average x value for each individual 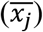. The within-individual effect is fitted as the deviation from the individual mean for a given observation 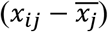 The previous equation becomes:

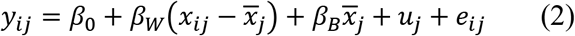

where *β_W_* is the slope of the within-individual effect and *β_B_* is the slope of the among-individual effect. Thus, equation 1 corresponds to a specific case of equation 2 when *β_W_* =*β_B_* and assumes that among- and within-individual effects are identical. van de Pol & Wright (2009) showed that a standard mixed model does not allow estimation of within-individual effects as reliably as the within-individual centering model.

## 2. The within-individual centering method to investigate quadratic patterns

The within-individual centering method has been regularly used in ecology to investigate quadratic changes in traits (11.5% of all the articles citing van de Pol & Wright (2009) and applying this method, Fig. 1). Quadratic terms could be added to both the within- and the among-individual effects independently since they can arise from different processes. Among the published articles using a quadratic relationship, all introduced it for the within-individual effect but only some of those used it simultaneously for the among-individual effect. The relevance of including quadratic terms to the within-individual effect, the among-individual effect, or both, depends on the system studied and the question investigated. Here, for the sake of clarity, we first present how quadratic terms can be added to the within- and among-individual effects separately. Then we give the general equation including a quadratic term for both within- and among-individual effects.

### 2.1 Quadratic within-individual effect

To use an individual centering approach including a quadratic within-individual effect, the individual centering should be applied on both linear and quadratic terms. We suggest the following equation to estimate within-individual quadratic effect (see Appendix S1 for full derivation and explanation of the equation):

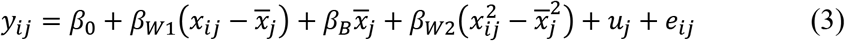

An alternative equation which has been used by most of the studies investigating quadratic individual trajectories is:

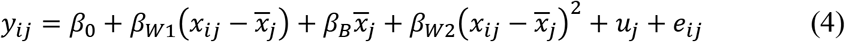

In equation 3, the quadratic within-individual effect (*β*_*W*2_) is included as the deviation of the squared *x* value from the squared average *x* value of an individual, i.e., 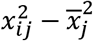, whereas in equation (4), the quadratic within-individual effect (*β*_*W*2_) is included as the square of the centered variable, i.e., 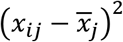. It should be noted that equations (3) and (4) are not equivalent and generate very different individual patterns (Fig. 3).

To better understand the difference between these two equations, we can develop equation (4) as:

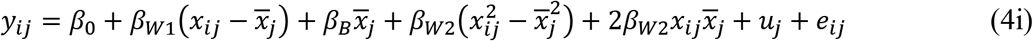

From equation (4i), we can see that the difference between the two equations is the presence of the term 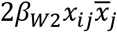 in equation (4) only. This term introduces an interaction between *x_ij_*, the value of a trait *x* at measurement *i* from individual *j*, and 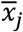, the average x value sampled for individual *j*. By doing so, equation (4) assumes that the quadratic within-individual effect (*β*_*W*2_) is not related to the absolute value of X but depends on the average 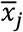 value sampled for each individual. Thus, equation (4) assumes that within-individual changes depends on the range of the explanatory variable sampled for each individual (Fig. 2B). In contrast, because there is no interaction between *x_ij_* and 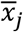 in equation (3), this equation assumes that the within-individual effect of X on Y depends on the absolute value of X. In other words, equation 3 assumes that there is a general quadratic pattern over the entire range of X which is shared by all the individuals. Ignoring individual variation due to the among-individual effect and random variation in the intercept, the effect of X on Y is the same for all individuals (Fig. 2A).

**Figure 2.**
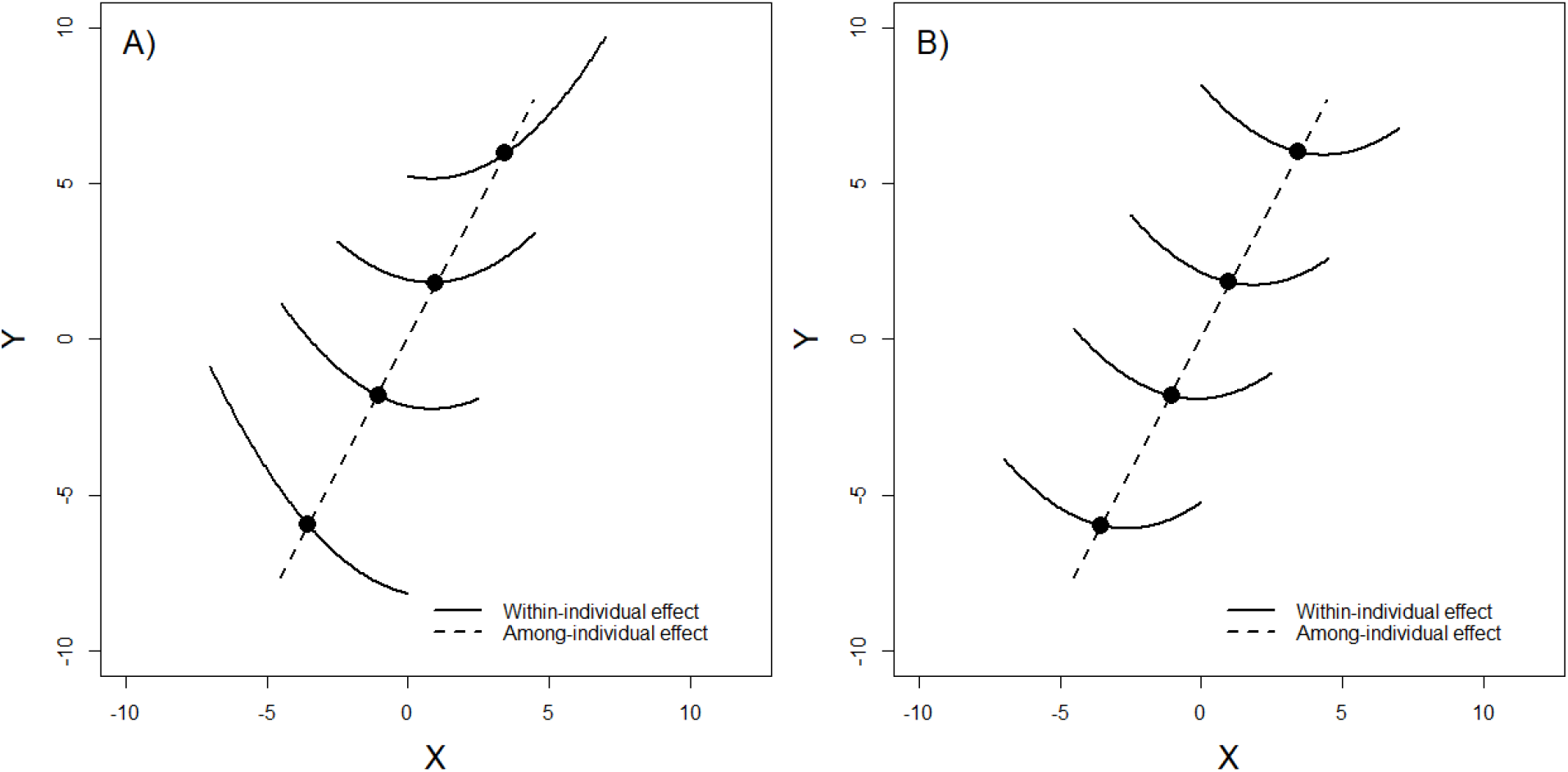
Individual trajectories illustrating quadratic within-individual effects simulated with equation (3) (A) and (4) (B). In both graphs, the four same individual life histories have been simulated using the equations (3) and (4), respectively, with the following parameters: *β*_0_ = 0, *β*_*W*1_ = −0.2, *β*_*W*2_ = 0.12, *β_B_*= 1, 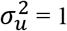 and 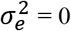. Black circles show 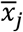, the average X values for each individual. The equation (3) (A) assumes that all individuals show the same pattern over the range of the explanatory variable. In contrast, equation (4) (B) assumes that the intra-individual changes of a given individual depend on the range of the explanatory variable sampled for this individual.

As a consequence, the two equations differ in where the quadratic pattern is expected to reach its maximum/minimum within each individual. Equation 3 assumes that the maximum/minimum is identical for all individuals on the absolute scale of X values, independently of the ranges of each individual trajectory. Equation 4 assumes that the maximum/minimum is different among individuals but is reached at the same relative place within the range of the explanatory variable observed for each individual (mean average measured for each individual, for instance). See appendix S2 for a mathematical demonstration of this difference.

### 2.2 Quadratic among-individual effect

To use an individual centering approach including a quadratic among-individual effect, equation (2) becomes:

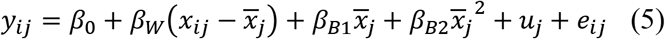

The quadratic among-individual effect (*β*_*B*2_) is fitted by the square of the average *x* value for each individual 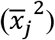 (Fig. 3). Contrary to the within-individual effect, we found no alternative equation in studies investigating quadratic individual trajectories. Because including a quadratic among-individual effect is not a source of confusion, the rest of the article focuses on the quadratic within-individual effect.

**Figure 3.**
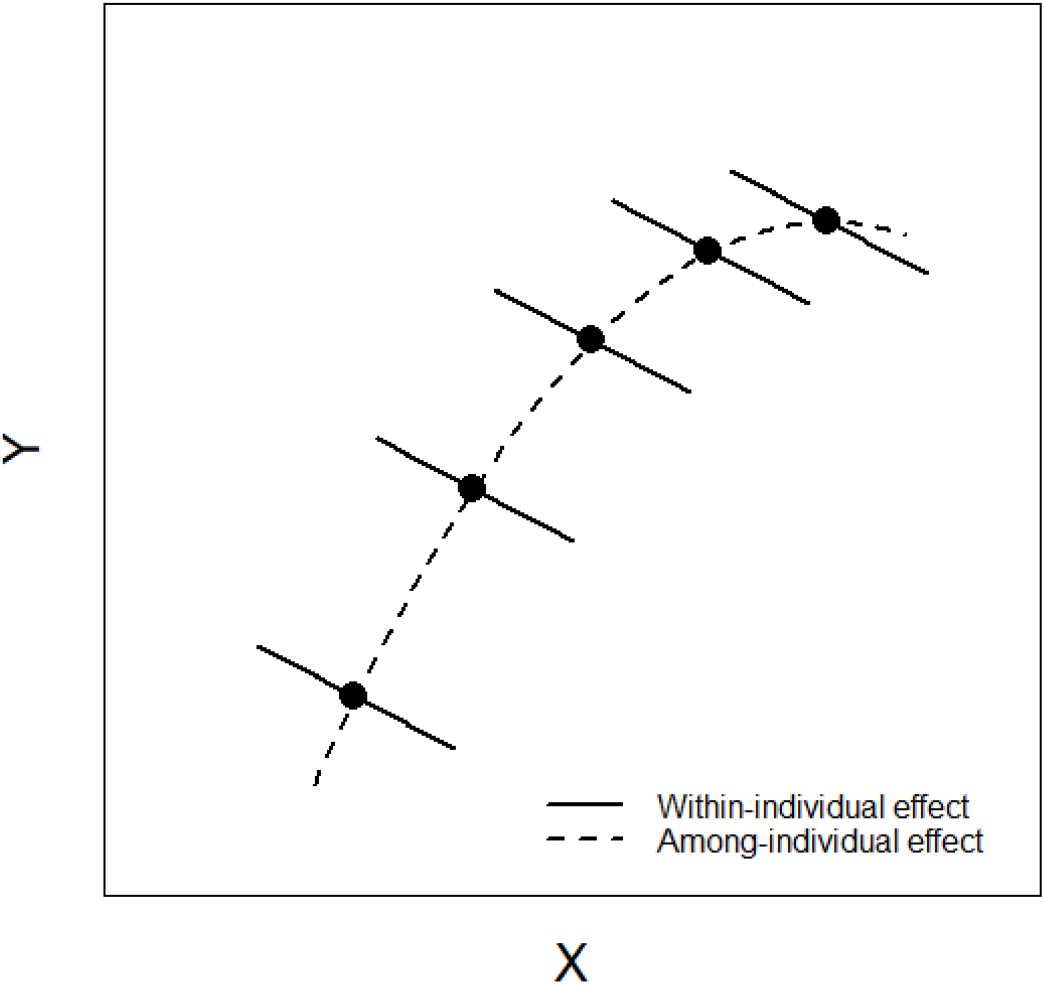
Individual trajectories illustrating a quadratic among-individual effect. Here, Y decreases linearly in relation to X within each individual but Y increases quadratically among individuals in relation to the average 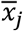 values. Individual trajectories are simulated with equation (5) with the following parameters: *β*_0_ = 0, *β_W_* = −0.5, *β*_*B*1_ = 1.2, *β*_*B*2_= −0.08, 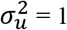 and 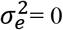. Black circles show 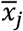.

### 2.3 Quadratic within-and among-individual effects

To use an individual centering approach including both a quadratic within-individual effect and a quadratic among-individual effect, the two alternative equations are:

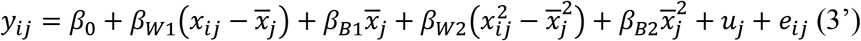

and

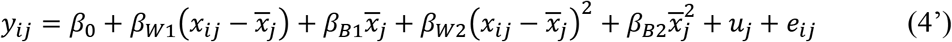

It should be noted that a key difference between the two approaches equations is that when within- and among-individual effects are similar, i.e., *β*_*W*1_ = *β*_*B*1_ and *β*_*W*2_ = *β*_*B*2_, only equation (3’) is equivalent to a model not using within-individual centering. Indeed, assuming identical within- and among-individual effect, we may simplify equation (3’) and (4’) as follows:

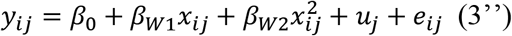

and

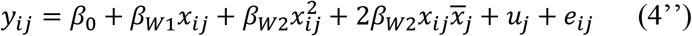

## 3. When should each equation be used? Examples of biological processes fitting each equation

Equation (3) assumes that all individuals show the same quadratic pattern over the range of the explanatory variable. Equation (3) can thus be used when researchers expect a functional relationship shared by all the individuals regardless of the mean value of the explanatory variable sampled for each individual (Figs. 3A & 6A). Taking the example of the flying speed performance in an insect, it is reasonable to expect consistent among-individual responses to variation in temperature. Because insects are ectotherms, flying performance of all individuals is expected to increase with temperature until a certain point where extreme temperature may have a negative effect. Here, there is a general quadratic pattern over the entire range of temperature shared by all the individuals (Fig. 4A). The effect of temperature on flying speed depends on the absolute value of the temperature, not on the deviation from the average temperature sampled for each individual. In this case, using equation (3) allows correct estimation of the quadratic within-individual effect. More generally, equation (3) could be used any time the within-individual response depends on the absolute value of the explanatory variable. This is expected to be the case in most studies investigating the response of individuals to biotic and abiotic environmental factors.

**Figure 4.**
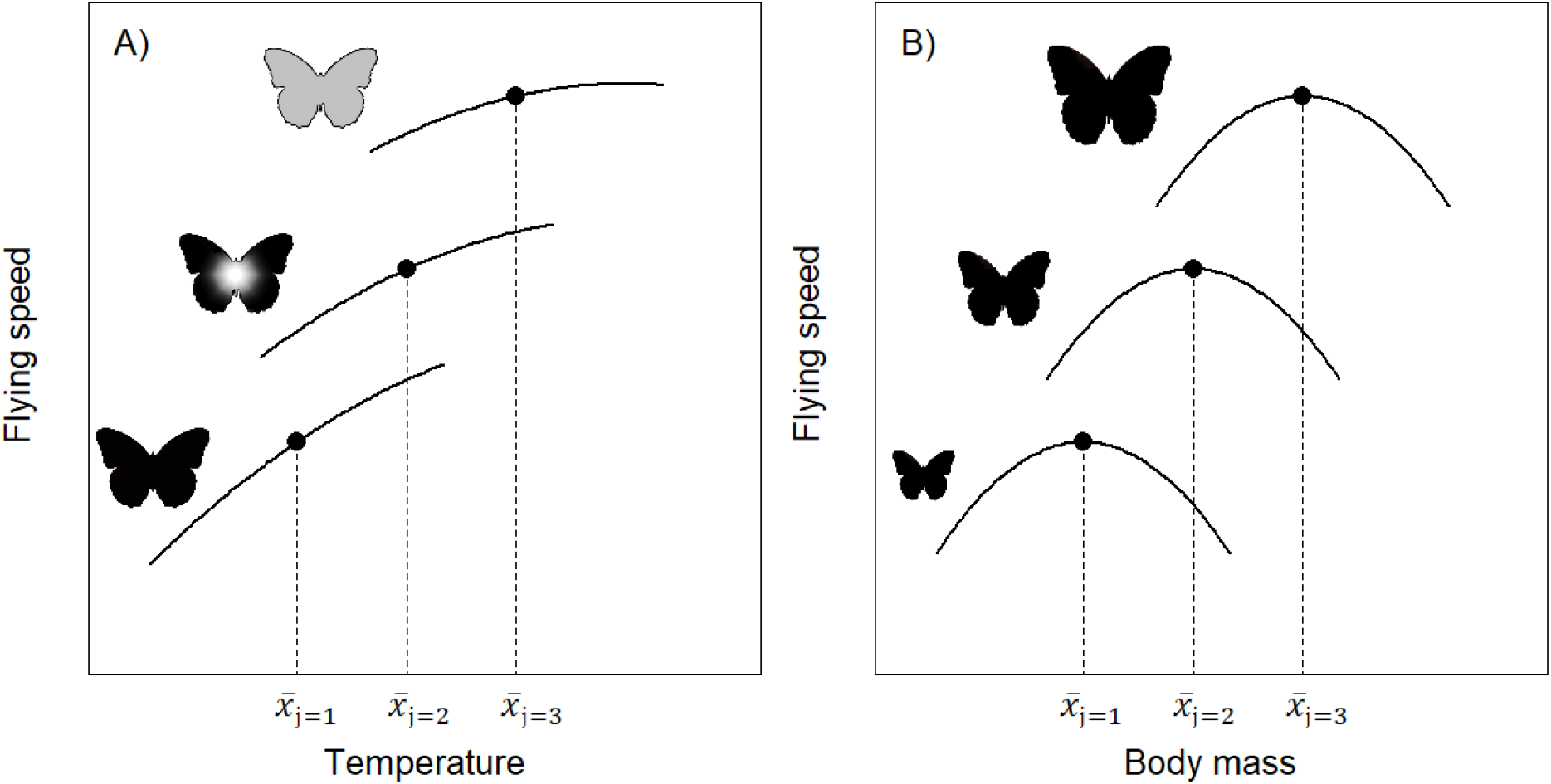
Theoretical effect of temperature (A) and body mass (B) on flying speed in a butterfly. These examples illustrate relationships following assumptions made by equation (3) and (4), respectively. Variation in flying speed is due to both a within-individual effect, i.e., effect of variation in temperature or body mass on individual performance, and an among-individual effect generated by variation in cohort (A) and size (B), for instance. In accordance with assumptions made by equation 3, all individuals show the same relationship between flying speed and temperature. In other words, the effect of the temperature on flying speed is independent of the range of temperature encountered by each individual. In contrast, variation in flying speed according to body mass shows a strong individually-specific response. Each quadratic pattern is centered on the mean body mass of each individual. In this case it is the mass deviation related to the individual optimum, not the absolute mass value, which is critical.

In contrast, equation (4) assumes that each individual has a quadratic pattern that depends on its individual mean. Continuing with the example of flying performance in an insect, an example of biological process that could fit equation 4 is the effect of body mass on flying performance. In this case, absolute body mass might not be critical to determine flying performance, but rather the deviation from the optimum individual body mass (Fig. 4B). As a consequence, patterns are centered on individual means in this example. The same change in body mass may be associated with higher or lower flying speed according to the size of the individuals. More generally, equation (4) assumes that the effect of the explanatory variable varies among individuals according to an individual optimum. This may correspond to any variables that have strong individual optima dependent on sampled individual *x_j_* values. It could for instance correspond to an acclimation process when individuals have optimal performance based on the environment they experienced in the past.

## 4. The case of quadratic age trajectories with selective disappearance: which equation should be used?

The importance in distinguishing within-individual from among-individual effects in age-related pattern has been recognized for a long-time (Forslund and Pärt 1995; Reid et al. 2003; Nussey et al. 2008). Unsurprisingly, the within-individual centering method has been commonly used in ecology to investigate age-related changes in behavioral, physiological, morphological or life-history traits (Fig. 1). When individual life-trajectories are considered in their entirety, the observed within-individual age patterns generally display a dome shape with increasing performance in early life and decreasing performance in late life due to senescence (Forslund and Pärt 1995; Aubry et al. 2009; Bouwhuis et al. 2009; Hämäläinen et al. 2014). Consequently, quadratic relationships have often been used to investigate age-related patterns. We found that 27% of the studies using the within-individual centering method to investigate age-related patterns included a quadratic within-individual effect. Among them, all studies describing the method with enough detail used equation (4), but none used equation (3). Although equation 4 has been applied systematically, this equation might be inappropriate for age-related patterns.

Using the within-individual centering method, the age effect is split into two terms: the average individual age whose associated slope estimates the among-individual effect (i.e., the effect due to selective disappearance of certain phenotypes), and the age-related changes (i.e., the difference between the actual and the average age of a given individual), whose slope estimates the within-individual effect. When we compare the individual-trajectories predicted by the equations (3) and (4), it appears that the equation 4 is not consistent with empirically observed age patterns (Loison et al. 1999; Reid et al. 2003; Zhang et al. 2015) (Fig. 5). For instance, equation 4 assumes that the age trajectory of an individual dying at 5 years old 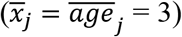 is similar to the age pattern between 6 and 10 years old of an individual dying at 15 years old 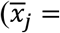 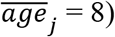. In both cases the age-related changes are the same, ranging from -2 to 2 (Fig. 4B). With equation (3), an individual dying young (low average X) is expected to display a trajectory that differs from that of an individual dying old (high average X). The former would show only the classical early-life increase, whereas the latter would show a more general pattern including an increase, a plateau and a decrease when older due to senescence (Fig. 5A). This pattern matches what is generally observed in longitudinal studies; that is, a decrease in individual performance occurring in later life after picking up in middle age (Loison et al. 1999; Reid et al. 2003; Zhang et al. 2015). In contrast, equation (4) does not allow modelling only the early-life increase for individuals dying young, but constrains all individuals to display the full quadratic pattern regardless of their age of death (Fig. 5B). This is generally inappropriate since in longitudinal studies, most of the individuals die before reaching the age of onset of senescence (Clutton-Brock 1988; Newton 1998). Finally, equation (4) makes the assumption of a fixed ageing pattern with a varying age of onset of senescence whereas equation (3) assumes a fixed age of onset of senescence estimating a general senescence pattern. It should be noted that extending equation 3 with random slopes for the linear and quadratic components of the within-individual effect would allow estimation of both varying rate and onset of senescence. However, those models, although theoretically sound, require much more data (van de Pol 2012; Westneat et al. 2020).

**Figure 5:**
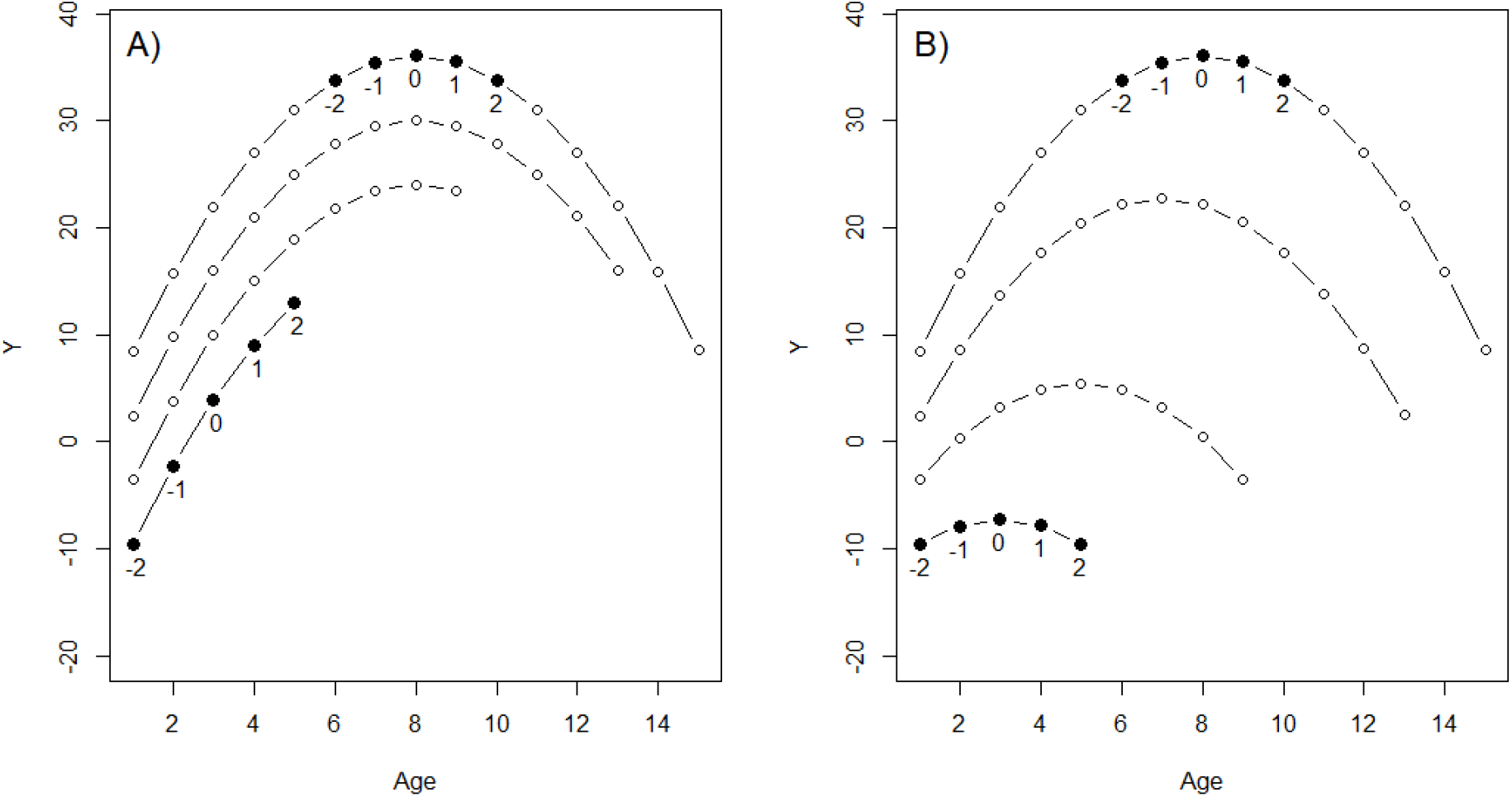
Age-related individual trajectories simulated according to equations 3 (A) and 4 (B). The five individual life histories are similar between the graphs in terms of the intercept and age range. Equation 4 assumes that the age pattern of an individual dying at 5 years old is similar to the age pattern between 6 and 10 years old of an individual dying at 15 years old (black dot with associated age-related changes).

In the case of age patterns, the among-individual effect may be generated by both selective appearance and disappearance effects (van de Pol and Verhulst 2006), making estimation of the among-individual effect more complicated. In addition, several studies strongly suggest moving away from polynomial models in favour of threshold models when studying aging patterns (Berman et al. 2009; Froy et al. 2017; Murgatroyd et al. 2018; Rodríguez-Muñoz et al. 2019). In this context, an interesting alternative method closely related to the within-individual centering is the use of age at first and last reproduction as fixed covariates (van de Pol and Verhulst 2006). This method allows disentangling of the selective appearance and disappearance, and facilitates the implementation of threshold models. For these reasons, we encourage the use of the method described by van de Pol and Verhulst (2006) instead of the more generic within-individual centering method in the case of age patterns.

## 5. Consequences of mismatches between generating and analytical equations

Above we showed that two alternative equations could be used for the investigation of quadratic within-individual effects. Here we illustrate the consequences of a mismatch between the equation generating the quadratic within-individual patterns and the equation used to analyze the data.

### 5.1. Methods

We simulated 1,000 data sets consisting of 50 individuals with 15 measurements each. First, for each individual, we used a normal distribution with a mean 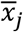, taken randomly between −7 and 7, and a variance 1.5 to sample 15 measurements. Then, we simulated the response variable Y using either equation (3) or (4). In both cases, we used the same parameters: *β*_0_ = 0, *β*_*W*1_ = 0.4, *β*_*W*2_ = −0.11, *β*_*B*1_= 1, *β*_*B*2_ =0, *σ_u_* = 2 and 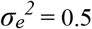. Finally, for each simulated dataset, we fitted three models: the matching and mismatching models (equation (3) and (4)) and a simpler linear model (equation 2).

Based on these sets of simulations, we assessed the consequences for model selection and quality of estimates of analyzing the data with the mismatching equation. We used the Akaike information criterion (AIC) to compare matching, mismatching and linear models. This comparison allows us to check if a mismatch between the generating and analytical equations affects our ability to detect quadratic within-individual effects. When differences of AIC between the two models were below 2, we followed the principle of parsimony, selecting the model with the lower number of parameters. To assess the effect of mismatches between generating and analytical equations on the quality of the estimates, we computed bias by subtracting the estimated parameters from the true parameter values used to generate the data. Results were summarized with violin plots to visualize both mean and dispersion of bias. Finally, we also computed the mean square error (MSE) which gives the accuracy of the estimates, i.e., combination of bias and precision to define the performance of an estimator.

### 5.2. Results

Quadratic models matching the equation used to simulate the data were always selected as the best of the three models (lower AIC). However, simpler linear models were selected against a mismatching quadratic model in 3% of the data simulated with equation (3) and analyzed with equation (4) and in 38% of the data simulated with equation (4) and analyzed with equation (3). This demonstrates that using the inappropriate equation may lead to flawed inferences.

Regarding the performance of the estimators, no bias was observed when the equation used to generate the data matched the one use to analyze the data (equation (3) bias *β*_*w*1_= −0.0005[−0.027:0.024] and *β*_*w*2_ = −0.0001[−0.003: 0.003], equation (4) bias *β*_*w*1_= −0.0009[− 0.027:0.025] and *β*_*w*2_ = −0.0001[−0.012: 0.013]) (Fig. 6). However, it should be noted that the performance of equation (4) depends on how accurately the average explanatory variable sampled for each individual corresponds to the true/biological average explanatory variable. When these quantities differ, for instance due to sampling bias, estimates of the quadratic within-individual pattern could be biased (Appendix S3).

**Figure 6:**
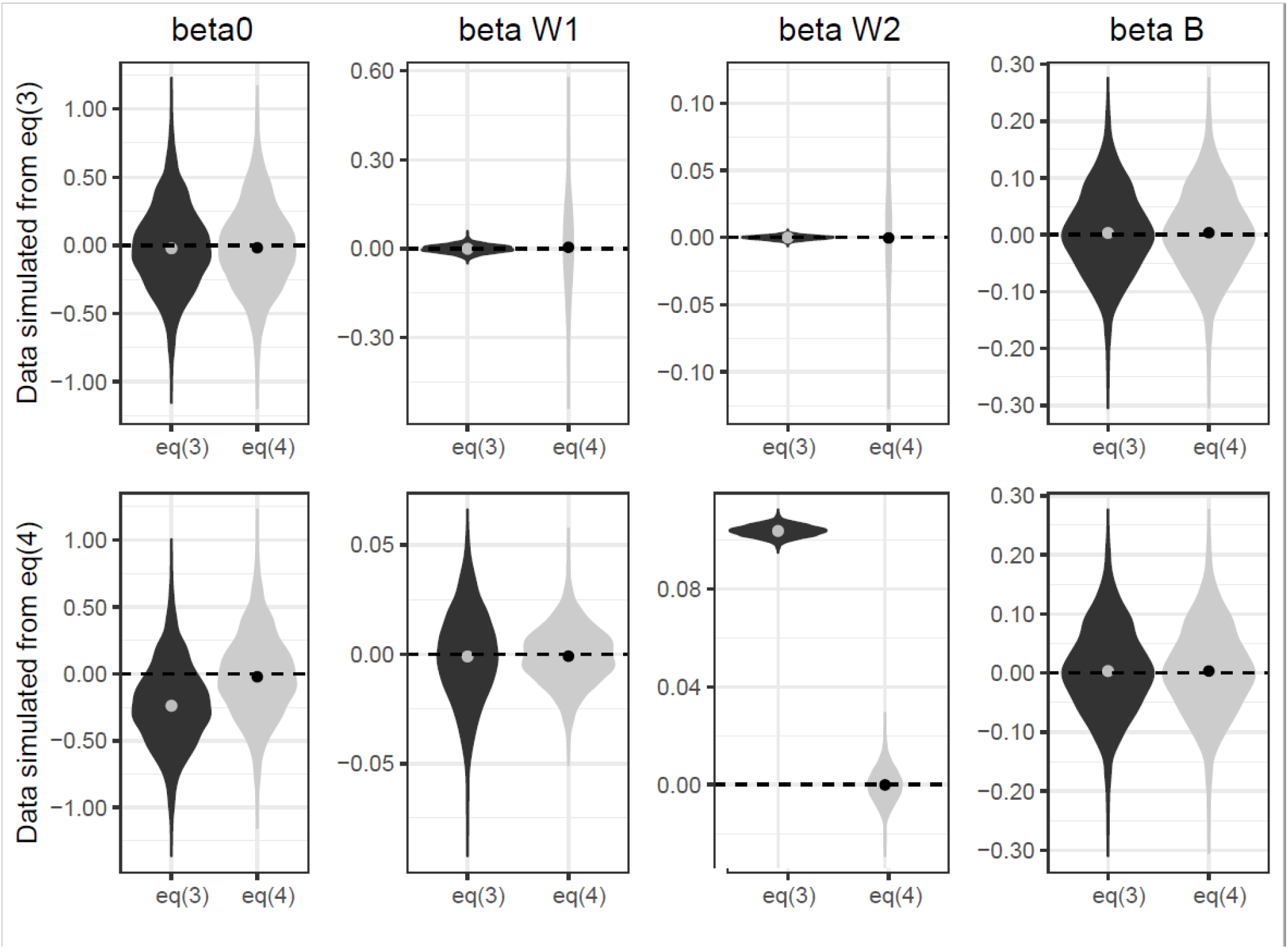
Bias in the estimates of the within- and among-individual effects from 1000 datasets simulated and analyzed by the two alternative quadratic equations of the within-centering method (data analyzed with equation 3 in black and equation 4 in grey). Parameters include the intercept (beta0), the linear (W1) and the quadratic (W2) within-individual effect of X, and the linear among-individual effect (B1).

Mismatch between the generating and analytical equations affects both bias and precision of the estimates. When data were simulated with equation 4 but analyzed with equation 3, the intercept was negatively biased and the quadratic within-individual effect was positively biased. When data were simulated with equation 3 but analyzed with equation 4, estimates were unbiased but very imprecise. In particular, the linear and quadratic within-individual estimates showed poor performance in this mismatching case (Figure 6 & S1). Whatever the mismatch, using the inappropriate equation may lead to strong misestimation of the within-individual pattern (Fig. 7). Nevertheless, the among-individual effect was estimated without bias and with a consistent precision whichever generating and analytical equations were used (Fig. 6).

**Figure 7:**
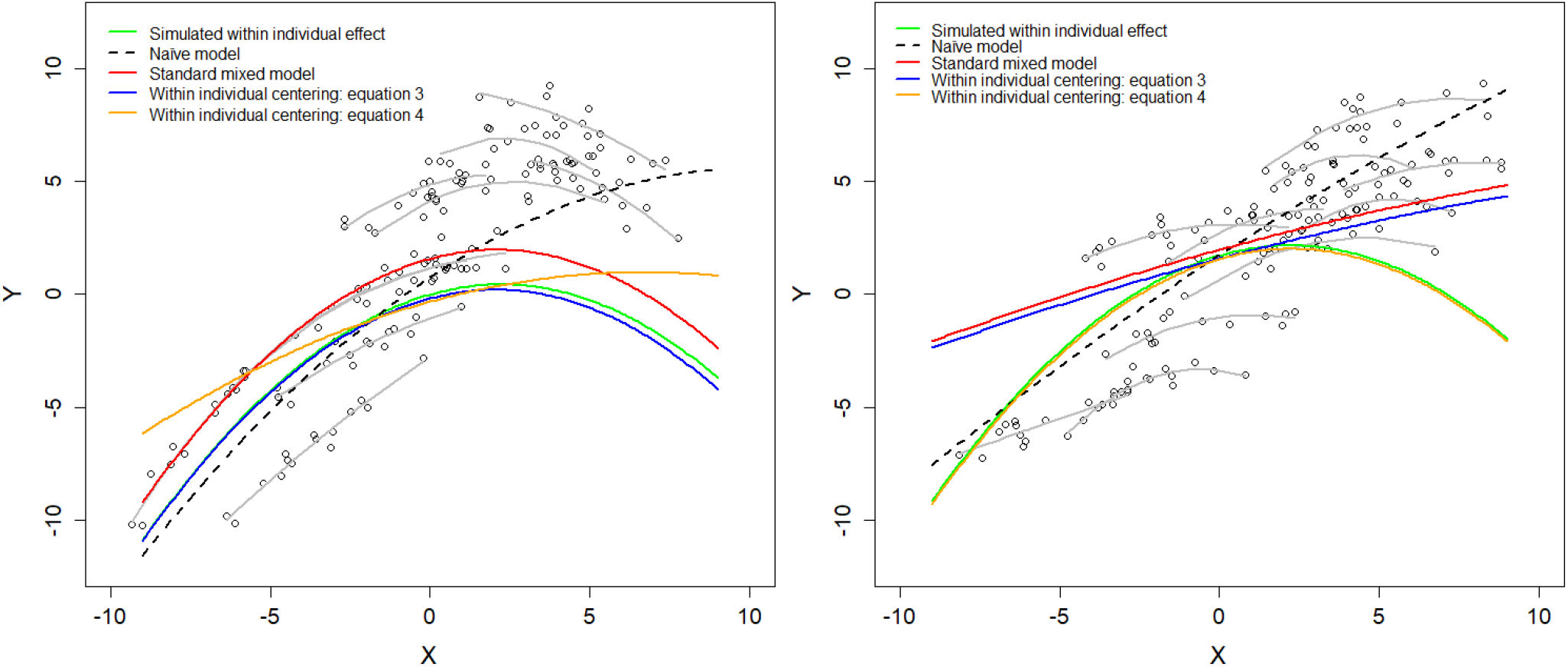
Estimation of the quadratic within-individual effect for data simulated with equation (3) (A) and (4) (B). Each dot represents one observation per individual and the gray lines depict observed within-individual trajectories. For illustration, we show the row observations for only 10 individuals. The green line shows the simulated within-individual effect, the dashed black line shows predictions of the naïve model that corresponds to a simple quadratic linear model, the red line shows predictions of the within-individual effect estimated using a standard quadratic mixed model (quadratic extension of equation 1), and the blue and orange lines show prediction of the within-individual effect estimated using the within-individual centering method according to equation 3 and 4 respectively.

## 6. Simulation of quadratic age trajectories with selective disappearance

### 6.1. Methods

We ran further simulations for age-related patterns corresponding to the most frequent individual-centered variable used for quadratic patterns. Instead of using equations (3) and (4), we generated age-related data with selective disappearance.

First, we simulated individual age-related trajectories using the following equation:

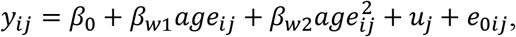

where *β*_0_ = 0 is the intercept, *β*_*w*1_ = 0.8 and *β*_*w*2_ = −0.06 are the linear and quadratic within-individual effects respectively, *u_j_* is deviation from the intercept for individual *j* assumed to be drawn from a normal distribution with zero mean and among-subject variance 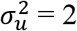, and *e_ij_* is the residual error assumed to be drawn from a normal distribution with zero mean and variance 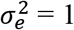. To simulate an among-individual effect, we generated the selective disappearance of frail individuals simulating age of death as a function of the individual-specific deviation in the response variable (*u_j_*). This differential survival among individuals in relation to their age-independent performance corresponds to what has been reported by empirical studies as individual quality (McCleery et al. 2008; Pigeon et al. 2017; Vedder and Bouwhuis 2018). We simulated individual age at death using an annual individual survival probability of ilogit(u_j_/2+1.5). Individuals with negative deviation, that is those with lower performances, also have lower survival prospects (see appendix S4 for the code used). Using the procedure described above, we simulated 1000 datasets with perfect detection, each consisting of 50 individuals. We analyzed these datasets assessing the relationships between the response variable and age with both equations (3) and (4) and computed bias and MSE as explained in section 5.1. Finally, because detection is often imperfect, we performed the same simulation analysis but with individual recapture probability set to 0.5 and to 0.2 instead of 1.

### 6.2. Results

Results confirmed that equation (3), but not equation (4), is suitable for the investigation of quadratic age patterns when selective disappearance occurs. Estimates of the within-individual age effect from equation (4) were strongly biased (bias *β*_*w*1_= −0.818[−0.860:−0.763] and *β*_*w*2_ = −0.0002[−0.008: 0.008]). Bias was higher in *β*_*w*1_ than in *β*_*w*2_ because *β*_*w*1_ allows counterbalancing of the supplemental component 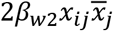 included in equation (4). Only equation (3) provided unbiased estimates (bias *β*_*w*1_ = −0.002[−0.093: 0.097] and *β*_*w*2_ = −0.0001[−0.006: 0.006]) (Figure S2). Using equation (4) instead of equation (3) for describing age-related individual trajectory led to flawed inferences with a strong overestimation of senescence (earlier onset and higher rate, Figure S3). Since real data often show incomplete histories of individual trajectories, i.e., detection probability is typically lower than 1, we reassessed the performance of these models considering incomplete individual capture histories. Results show that missing information within the lifetime of an individual does not change the respective performance of the models previously considered, but increases uncertainty for all parameters. Even when the detection probability is very low i.e., 0.2, leading to very patchy capture histories for each individual, the within-individual centering estimates from equation (3) are accurate whereas those from equation (4) are consistently biased (Figure S2 & S4).

## Conclusion

The within-individual centering method should be used with caution for non-linear relationships. We showed that two different equations could be used to estimate within-individual quadratic relationships and that each equation makes different assumptions about the shape of the individual pattern. The choice of using one equation instead of the other should depend upon the biological process investigated. We believe that equation (3), which assumes that the individual response depends on the absolute value of the explanatory variable, better fits many biological processes than equation (4), which assume that the individual response depends on relative value of the explanatory variable to the subject mean. Typically, poor and good conditions in climatic factors, food abundance or parasite load is expected to be uniform among-individuals as assumed by equation 3 (Fig. 3A). Equation (4) assumes that the average explanatory variable sampled for each individual matches the average condition driving the quadratic within-individual pattern. It is questionable whether this assumption is frequently met in practice. In many ecological studies, the range of environmental conditions sampled for a given individual is at least in part randomly determined. For instance, if we investigate the individual response to a climatic variable such as precipitation, the range of values considered for each individual will depend on the conditions occurring during the period of observation of those individuals in the field. As a consequence, even if the process generating the response variable corresponds to equation 4, the average explanatory variable sampled for each individual would not necessarily match the average condition driving the quadratic within-individual pattern.

Using the within-individual centering approach with polynomial terms requires larger sample sizes at the individual level to be able to properly assess the linear and quadratic patterns both at the within- and among-individual levels. Even if the simulations showed no biases when matching equations were used, it should be noted that we used data sets with numerous repeated measures (50 individuals with 15 observations each). The biases and reliability of within-individual centering with quadratic patterns have not yet been properly assessed and we urge researchers to consider their sample size and data structure before using this type of approach.

In any case, when using the within-individual centering for non-linear relationships, the equation used should be chosen carefully in order to run meaningful analyses. In line with Westneat et al (2020), we stress that it is critical to have a clear understanding of the equation used to ensure that the assumptions made by a statistical model (using within-individual centering or not) match the biological process investigated. We hope that this manuscript will encourage the use of the within-individual centering method and promote its correct application for non-linear relationships.

## Acknowledgments

The authors are particularly grateful to M. Authier and S. Robin for stimulating discussions and to J-F Lemaître, M. van de Pol, D. Nussey, J-M. Gaillard and D. Westneat and an anonymous reviewer for valuable comments on the manuscript. Finally, we thank A. Nicol-Harper for correcting our English.

