## Appendix for "Distinguishing within- from between-individual effects: How to use the within-individual centering method for quadratic patterns"

**Limits of the within centering method**

**Supplementary material**

Appendix S1: Justification and explanation of equations 3 and 3’ in the main text.

In this manuscript, we suggest the use of $\overline{x}_{j}^{2}$ instead of $\overline{x_{j}^{2}}$ to center the squared variable *x*. We explain below the details of our reasoning.

The core idea of the within-individual centering approach is to separate the effect of an explanatory variable in two parts: 1) the among-individual effects due to difference in mean explanatory variable experienced by the different individuals and 2) the within-individual effect that reflects the individual response to the explanatory variable. To implement a within-individual centering approach on a polynomial equation, the within-individual centering should be done on each term independently. Thus, we start with the equation of $y$ as a function of a linear and quadratic terms of $x$ for observations $i$ and individual $j$

$y_{ij}=\beta_{0}+\beta_{1}x_{ij}+\beta_{2}x_{ij}^{2}+u_{j}+e_{ij}$ (S1)

where $u_{j}$ is the deviation from the intercept for individual $j$ and $u_{j}\sim N\left( 0,\sigma_{id}^{2} \right)$

If we want to use the within-individual centering approach, we need to apply it on both linear and quadratic terms.

$y_{ij}=\beta_{0}+\beta_{w_{1}}\left( x_{ij}-\overline{x_{j}} \right)+\beta_{b1}\overline{x_{j}}+\beta_{w2}\left( x_{ij}^{2}-\overline{x_{j}^{2}} \right)+\beta_{b2}\overline{x_{j}^{2}}+u_{j}+e_{ij}$	(S2)

where $\beta_{w_{1,2}}$ and $\beta_{b_{1,2}}$ are the within- and among-individual effects for the linear and quadratic terms.

However, one major problem with this approach is that $\overline{x_{j}^{2}}$ is proportional to the variance in $x_{j}$ since $\sigma_{x_{j}}^{2}=\overline{x_{j}^{2}}-{\overline{x_{j}}}^{2}$. This approach would lead to biases when the variance in $x$ differs among individuals. To show why, we can introduce the variance terms in equation S2. We get:

$y_{ij}=\beta_{0}+\beta_{w_{1}}\left( x_{ij}-\overline{x_{j}} \right)+\beta_{b_{1}}\overline{x_{j}}+\beta_{w_{2}}\left( x_{ij}^{2}-\sigma_{x_{j}}^{2}-{\overline{x_{j}}}^{2} \right)+\beta_{b_{2}}\left( \sigma_{x_{j}}^{2}+{\overline{x_{j}}}^{2} \right)+u_{j}+e_{ij}$ (S3)

which gives, after rearranging:

$y_{ij}=\beta_{0}+\beta_{w_{1}}\left( x_{ij}-\overline{x_{j}} \right)+\beta_{b_{1}}\overline{x_{j}}+\beta_{w_{2}}\left( x_{ij}^{2}-{\overline{x_{j}}}^{2} \right)+\beta_{b_{2}}{\overline{x_{j}}}^{2}+\left( \beta_{b_{2}}-\beta_{w_{2}} \right)\sigma_{x_{j}}^{2}+u_{j}+e_{ij}$ (S4)

We can see that when $\beta_{b_{2}}=\beta_{w_{2}}$, that is when among-individual and within-individual quadratic effects are similar, the variance in $x_{j}$ has no impact on $y_{ij}$ since the term $\left( \beta_{b_{2}}-\beta_{w_{2}} \right)\sigma_{x_{j}}^{2}$ equals 0. However, as it is expected that among- and within-individual effects differ in most cases, using equation S2 (equivalent to S4) could introduce biases because the term $\left( \beta_{b_{2}}-\beta_{w_{2}} \right)\sigma_{x_{j}}^{2}$ is generally not equal to 0. Assuming that the data is generated according to equation S2, but the sampling has been done on a restricted part of the parameter space and thus that the sampled variance in $x$ for individual $j$ ($V_{x_{j}}$) is different from the variance in $x$ for individual $j$ in the generating process $\sigma_{x_{j}}^{2}$, then the estimated values of $\beta_{b_{2}}$ and $\beta_{w_{2}}$ could be strongly biased. In addition, using equation S2 would make comparison of parameter estimates harder across studies when the variance in x for each individual differs across individuals and studies. Finally, from a data generation perspective, equation S2 would predict different responses for individuals with equal $x_{j}$ but different variance in $x_{j}$.

Alternatively, based on equation S4, we can see that using $\overline{x}^{2}$ instead of $\overline{x^{2}}$ to center the quadratic effect at the within-individual level should provide similar estimates of $\beta_{b_{1,2}}$ and $\beta_{w_{1,2}}$. We can thus fit a model using a within-individual centering approach with a quadratic relation using:

$y_{ij}=a+b_{w_{1}}\left( x_{ij}-\overline{x_{j}} \right)+b_{b_{1}}\overline{x_{j}}+b_{w_{2}}\left( x_{ij}^{2}-{\overline{x_{j}}}^{2} \right)+b_{b_{2}}{\overline{x_{j}}}^{2}+id_{j}+e_{ij}$	(S5)

which corresponds to equation (3’) in the main text. However, because $V_{x_{j}}$ is not explicitly accounted for, it would affect the estimate of the population intercept and it would increase the variance of the random effect individual identity $V_{id}$.

When $V_{x_{j}}$ is the same for all individuals then $\left( \beta_{b_{2}}-\beta_{w_{2}} \right)V_{x_{j}}$ will be the same for all individuals and should be captured in the estimate of the population intercept, thus $a=\beta_{0}+\left( \beta_{b_{2}}-\beta_{w_{2}} \right)V_{x_{j}}$

When $V_{x_{j}}$ differs among individuals then $\left( \beta_{b_{2}}-\beta_{w_{2}} \right)V_{x_{j}}$ will be split across the intercept $a$ and the random effect $u_{j}$ where $a=\beta_{0}+\left( \beta_{b_{2}}-\beta_{w_{2}} \right)\overline{V_{x_{j}}}$ and $id_{j}=u_{j}+\left( \beta_{b_{2}}-\beta_{w_{2}} \right)\left( V_{x_{j}}-\overline{V_{x_{j}}} \right)$ with $V_{id}=\sigma_{id}^{2}+\left( \beta_{b_{2}}-\beta_{w_{2}} \right)^{2}V_{V_{x_{j}}}$

Thus, when using the within-individual centering method for quadratic patterns with equation S5 (or 3’), it should be recognized that if the underlying generating equation is equation S1 or S2 then the among-individual variation in the intercept will be positively biased upward by a factor of the square of the difference between the among- and within-individual quadratic effect multiplied by the variance in the individual variance in $x$.

Appendix S2: Estimating the value at the maximum/minimum of the quadratic effect for equations 3’ and 4’

To estimate the value of $x$ at the maximum/minimum of $f\left( x \right)$, a quadratic polynomial of $x$, we need to solve the equation where the first derivative of $f\left( x \right)$ is equal to zero, $\frac{\delta f}{dx}=0$. Starting with equation 3’ from the main text (or S5 in appendix), we have:

$f\left( x,j \right)=\beta_{0}+\beta_{w_{1}}\left( x-\overline{x_{j}} \right)+\beta_{b_{1}}\overline{x_{j}}+\beta_{w_{2}}\left( x^{2}-{\overline{x_{j}}}^{2} \right)+\beta_{b_{2}}{\overline{x_{j}}}^{2}+u_{j}$	(S6)

For an individual $j$ $\overline{x_{j}}$ and ^2 are constant, so we get:

$\frac{\delta f}{dx}=\beta_{w_{1}}+2\beta_{w_{2}}x$	(S7)

solving for $\frac{\delta f}{dx}=0$, we obtain:

$x_{max/min_{j}}=\frac{-\beta_{w_{1}}}{2\beta_{w_{2}}}$	(S8)

For equation 3’, the value of $x$ at the maximum/minimum will be similar for all individuals since it is independent from individual variation in intercept, mean or variance of $x$ and will be located at $x=\frac{-\beta_{w_{1}}}{2\beta_{w_{2}}}$.

For equation 4’, we can do the same exercise. As a reminder equation 4’ is:

$g\left( x,j \right)=\beta_{0}+\beta_{w_{1}}\left( x-\overline{x_{j}} \right)+\beta_{b_{1}}\overline{x_{j}}+\beta_{w_{2}}\left( x-\overline{x_{j}} \right)^{2}+\beta_{b_{2}}{\overline{x_{j}}}^{2}+u_{j}$	(S9)

Taking the first order derivative of $g\left( x,j \right)$ for an individual $j$, we get:

$\frac{\delta g}{dx}=\beta_{w_{1}}+2\beta_{w_{2}}x-2\beta_{w_{2}}\overline{x_{j}}$	(S10)

solving for $\frac{\delta g}{dx}=0$, we obtain:

$x_{max/min_{j}}=\frac{{-\beta}_{w_{1}}}{2\beta_{w_{2}}}+\overline{x_{j}}$	(S11)

For equation 4’, the value of $x$ at the maximum/minimum can be different according to the individuals since it is dependent of the individual mean value of x.

Appendix S3: Bias in the estimates of the within- and among-individual effects for quadratic pattern according to equation 4, when the sampled individual mean $\overline{x}_{j}$ is different from the true individual mean $\overline{X}_{j}$.

Here we investigate the consequence when the sampled individual mean $\overline{x}_{j}$ is different from the true individual mean $\overline{X}_{j}$. We simulated 1000 datasets each consisting of 50 individuals for which we have 15 measurements. For each individual, we randomly simulated an individual mean $\overline{X}_{j}$ between -7 and 7. Then, instead of using a normal distribution centered on the $\overline{X}_{j}$, we used a uniform distribution between -7 and 7 to sample 15 measurements per individual. Thus, the 15 measurements are sampled independently from the location of $\overline{X}_{j}$ and consequently the sampled individual mean $\overline{x}_{j}$ differs from $\overline{X}_{j}$. Then, we generated the response variable Y using equation (4) and the individual mean $\overline{X}_{j}$. For the parameters, we used $\beta_{0}$ = 0, $\beta_{W1}$ = 0.4, $\beta_{W2}$ = -0.11, $\beta_{B1}$= 1, $\beta_{B2}$= 0, *σ_u_* = 2 and *σ_e_^2^* = 0.5. Finally, we analyzed the relationships between sampled explanatory and response variables using the equation 4. For this analysis, we use the sampled individual mean $\overline{x}_{j}$ which differ to some degree from $\overline{X}_{j}$ which has been used to generate the data. Bias and mean squared errors are presented in the following figure.

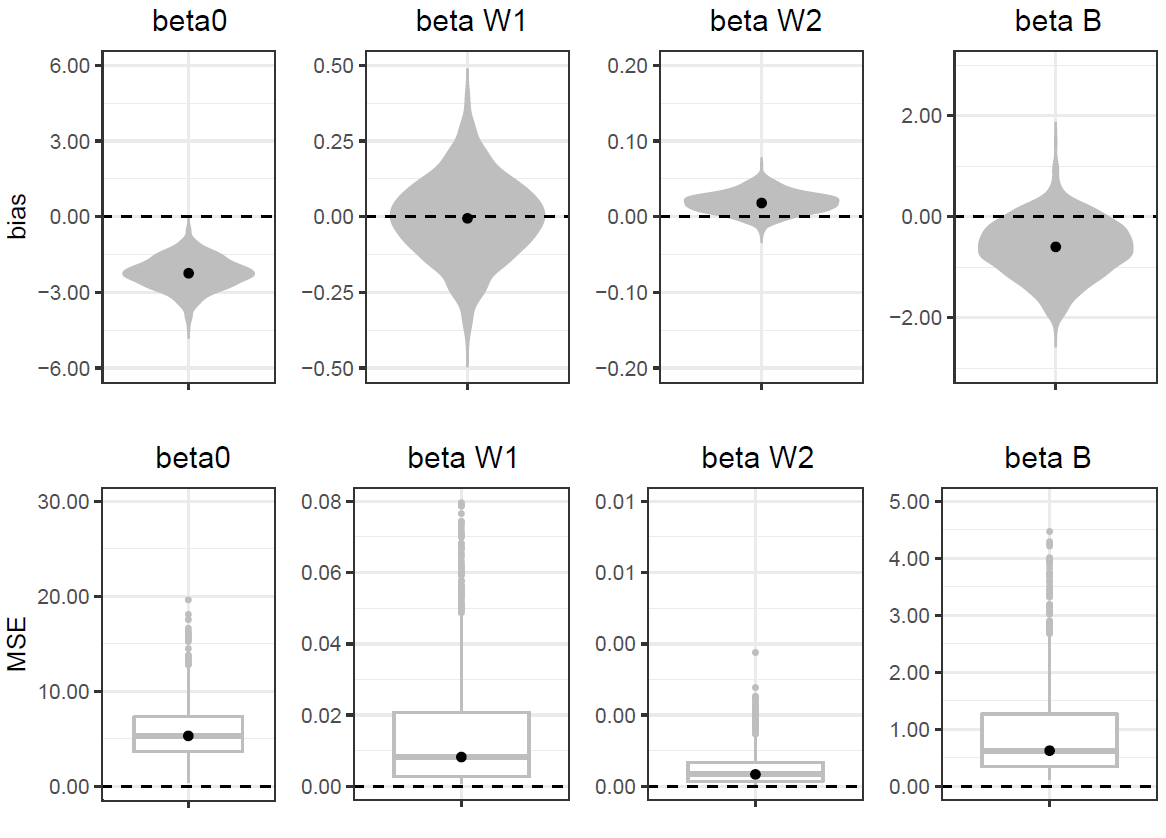

Appendix S4: Code used to simulate a quadratic age-dependence with selective disappearance.

```
ilogit <- function(x){1/(1+exp(-x))}

n.ind <- 500
n.year <- 16
beta0 <- 0
sigma.u <- 2
sigma.e <- 1
BW1 <- 0.80
BW2 <- -0.06
precap <- 1

## Simulation of individual age trajectories
y.mat <- matrix(data = NA, nrow = n.ind, ncol = n.year)
u <- array()
for (i in 1:n.ind){
  u[i] <- rnorm(1, 0, sigma.u)
  for (j in 1:n.year){
    e <- rnorm(1, 0, sigma.e)
    y.mat[i,j] <- beta0 + BW1 * (j-1) + BW2 * (j-1)^2 + u[i] + e
  }
}

# Simulation of survival data
s.mat <- matrix(data = 1, nrow = n.ind, ncol = n.year)

for (i in 1:n.ind){
  for (j in 2:n.year){
    s.mat[i,j] <- rbinom(1,1,ilogit(u[i]/2+1.5))*s.mat[i,j-1]
  }
}

#capture probability
data.Sp <- s.mat
for (i in 1:n.ind){
  for (j in 2:n.year){
    data.Sp[i,j] <- rbinom(1,1,precap)*s.mat[i,j]
  }
}

# Computation of new age-metric for the within individual centering
mean.age <- mean.age2 <- last.capture <- array(NA)
for (i in 1:n.ind){
  age.max <- max(which(data.Sp[i,]==1))
  mean.age[i] <- mean(c(1:age.max))
  mean.age2[i] <- mean(c(1:age.max)^2)
  last.capture[i] <- age.max
}
age <- matrix(data = rep(1:n.year,each=n.ind), nrow = n.ind, ncol = n.year)* data.Sp
```

Figure S1: Mean squared error in the estimates of the within- and among-individual effects from 1000 datasets simulated and analyzed by the two alternative quadratic equations of the within-centering method (equation 3 in black and equation 4 in grey). Parameters include the intercept (beta0), the linear (W1) and the quadratic (W2) within-individual effect of X, and the linear among-individual effect (B1).

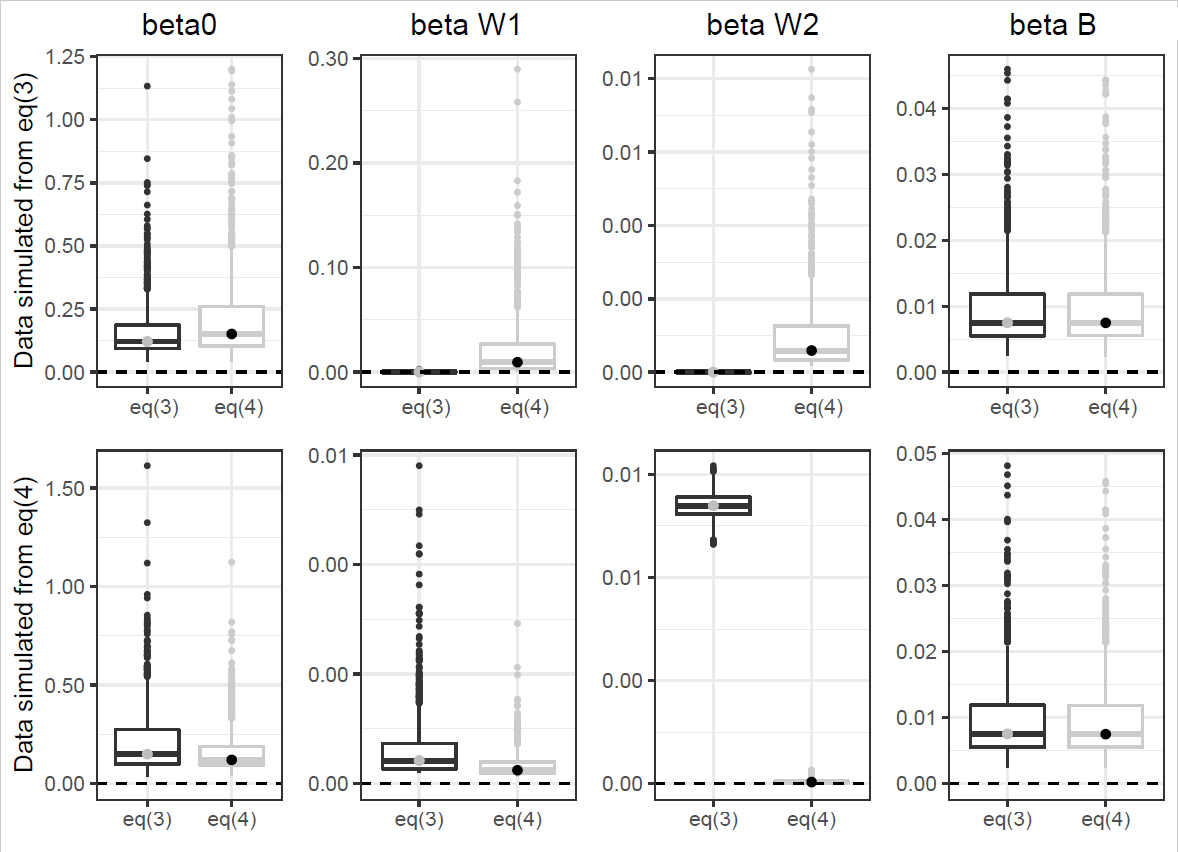

Figure S2: Bias in model estimates in the case of quadratic age-dependence with selective disappearance.

In addition to equations 3 and 4, we estimated the within-individual effect using the *naive model (naive)* that corresponds to a simple regression providing the average trajectory in the population, the *standard mixed model* (stand., quadratic version of equation 1 in the main manuscript) and the method described by van de Pol and Verhulst (2006, *2006*). Parameters include the intercept (beta0), the linear within-individual effect of age (beta W1) and the quadratic within-individual effect of age (beta W2). The following graphs show the distribution of the bias based on 1000 simulated datasets. Simulations have been done considering both complete individual life-history (recapture probability of one) and incomplete individual life-history (recapture probability of 0.5 and 0.2).

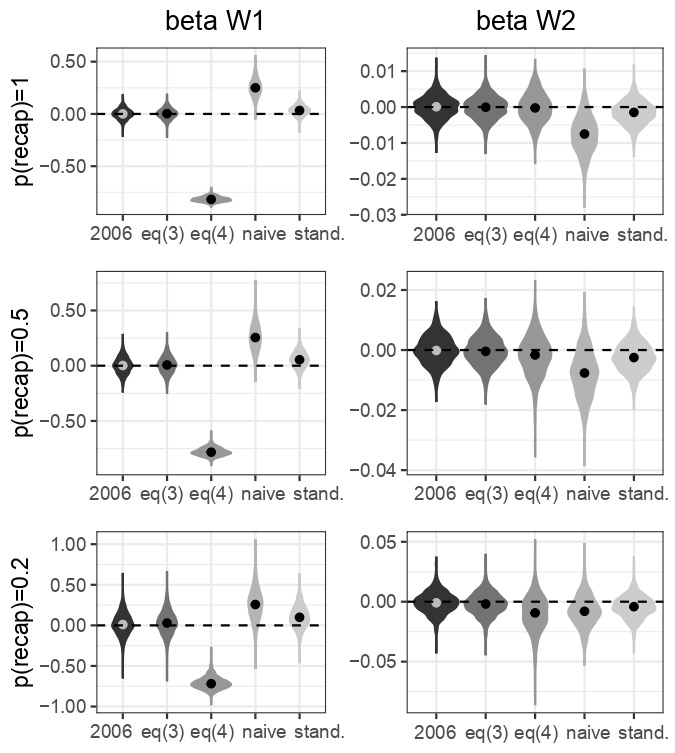

Figure S3: Estimation of the quadratic within-individual effect for data simulated with selective disappearance. The green line shows the simulated within-individual effect, the dashed black line shows the predictions of the naïve model that corresponds to a simple quadratic linear model, the red line shows the predictions of the within-individual effect estimated using a standard mixed model, and the blue and orange lines show the predictions of the within-individual effect estimated using the within-individual centering method according to equations 3 and 4 respectively. Each dot is one observation per individual and the gray lines depict observed within-individual trajectories. For illustration, we show the row observations for only 10 individuals.

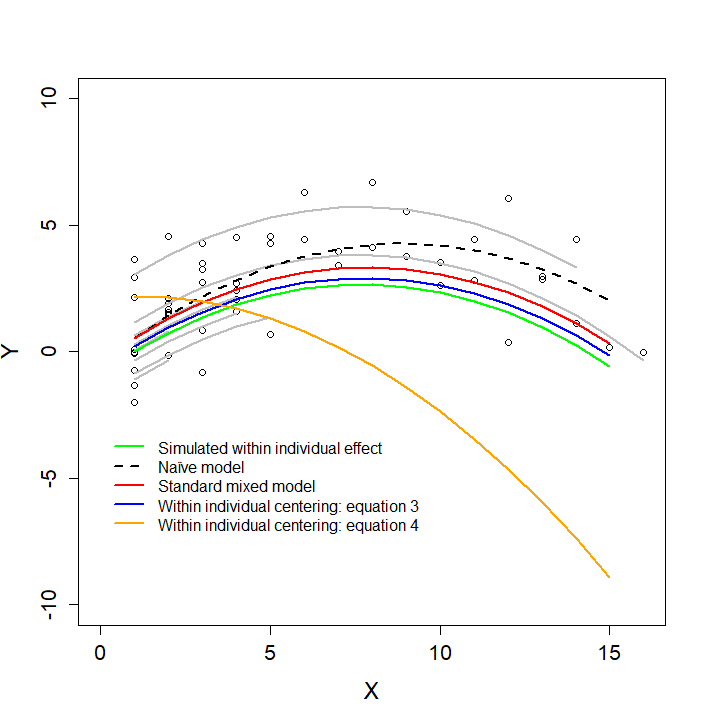

Figure S4: Mean squared error in model estimates in the case of quadratic age-dependence with selective disappearance.

In addition to equations 3 and 4, we estimated the within-individual effect using the *naive model (naive)* that corresponds to a simple regression providing the average trajectory in the population, the *standard mixed model* (stand., quadratic version of equation 1 in the main manuscript) and the method described by van de Pol and Verhulst (2006, *2006*). Parameters include the intercept (beta0), the linear within-individual effect of age (beta W1) and the quadratic within-individual effect of age (beta W2). The following graphs show the distribution of the bias based on 1000 replicates. Simulations have been done considering both complete individual life-history (recapture probability of one) and incomplete individual life-history (recapture probability of 0.5 and 0.2).

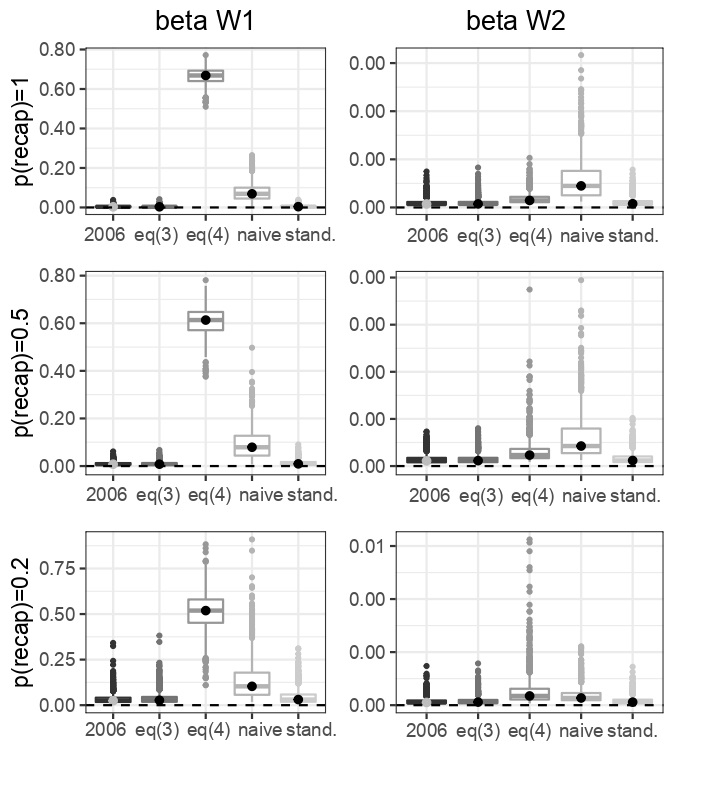
